# The macaque anterior cingulate cortex translates counterfactual choice value into actual behavioral change

**DOI:** 10.1101/336917

**Authors:** E Fouragnan, BKH Chau, D Folloni, N Kolling, L Verhagen, Miriam Klein-Flügge, L Tankelevitch, GK Papageorgiou, JF Aubry, J Sallet, MFS Rushworth

**Author notes:** Authors contributed equally to the work (E Fouragnan, BKH Chau, D Folloni). Authors contributed equally to the work (J Sallet, MFS Rushworth). To whom correspondence should be addressed: Address: Elsa Fouragnan, School of Psychology, University of Plymouth, Drake Circus, Plymouth, PL4 8AA, UK. Correspondence for the Transcranial Focused Ultrasound Stimulation technique: Address: Jean-Francois Aubry, Institut Langevin, Espci Paris, CNRS UMR 7587, INSERM U979 17 rue Moreau, Paris 75012, France.

## Abstract

The neural mechanisms mediating sensory-guided decision making have received considerable attention but animals often pursue behaviors for which there is currently no sensory evidence. Such behaviors are guided by internal representations of choice values that have to be maintained even when these choices are unavailable. We investigated how four macaque monkeys maintained representations of the value of counterfactual choices – choices that could not be taken at the current moment but which could be taken in the future. Using functional magnetic resonance imaging, we found two different patterns of activity co-varying with values of counterfactual choices in a circuit spanning hippocampus, anterior lateral prefrontal cortex, and anterior cingulate cortex (ACC). ACC activity also reflected whether the internal value representations would be translated into actual behavioral change. To establish the causal importance of ACC for this translation process, we used a novel technique, Transcranial Focused Ultrasound Stimulation, to reversibly disrupt ACC activity.

## Introduction

Every day, chacma baboons, an old world primate, navigate in their environment from the safety of their sleeping post, such as a cliff-top, to distant foraging or watering sites^1^. The decision to move towards one of those high valued locations is not simply guided by the accumulation of sensory evidence for that choice but by an internal representation or memory of the choices’ value. The same is true when they move back towards the sleeping post in the evening. While sensory and associative decision making have been well-studied^2^, we still lack an understanding of learning and decision making when currently unavailable options need to be held in memory and how those representations interact with current choices to guide behavior. Here we introduce a simple paradigm allowing investigation of the neural mechanisms mediating future decisions guided by information held in memory in macaques. In addition, the paradigm allows examination of how information held in memory modulates current on-going decisions.

Four macaques were trained to choose between pairs of abstract visual stimuli while in the MRI scanner (fig.1a, b). On each trial, the two stimuli available for choice (available options) were drawn from a set of three that were each associated with distinct reward probabilities (fig.1a). The probabilities drifted over trials and each stimulus’ reward probability was uncorrelated from that of the others (<22% mean shared variance). On each trial one of the two available options was chosen by the monkey, the other was unchosen, and a third option was invisible and unavailable for choice. Both the unchosen option and the unavailable option can be considered counterfactual choices – although these choices were not made on the current trial, they might be made on a future occasion.

**Figure 1.**
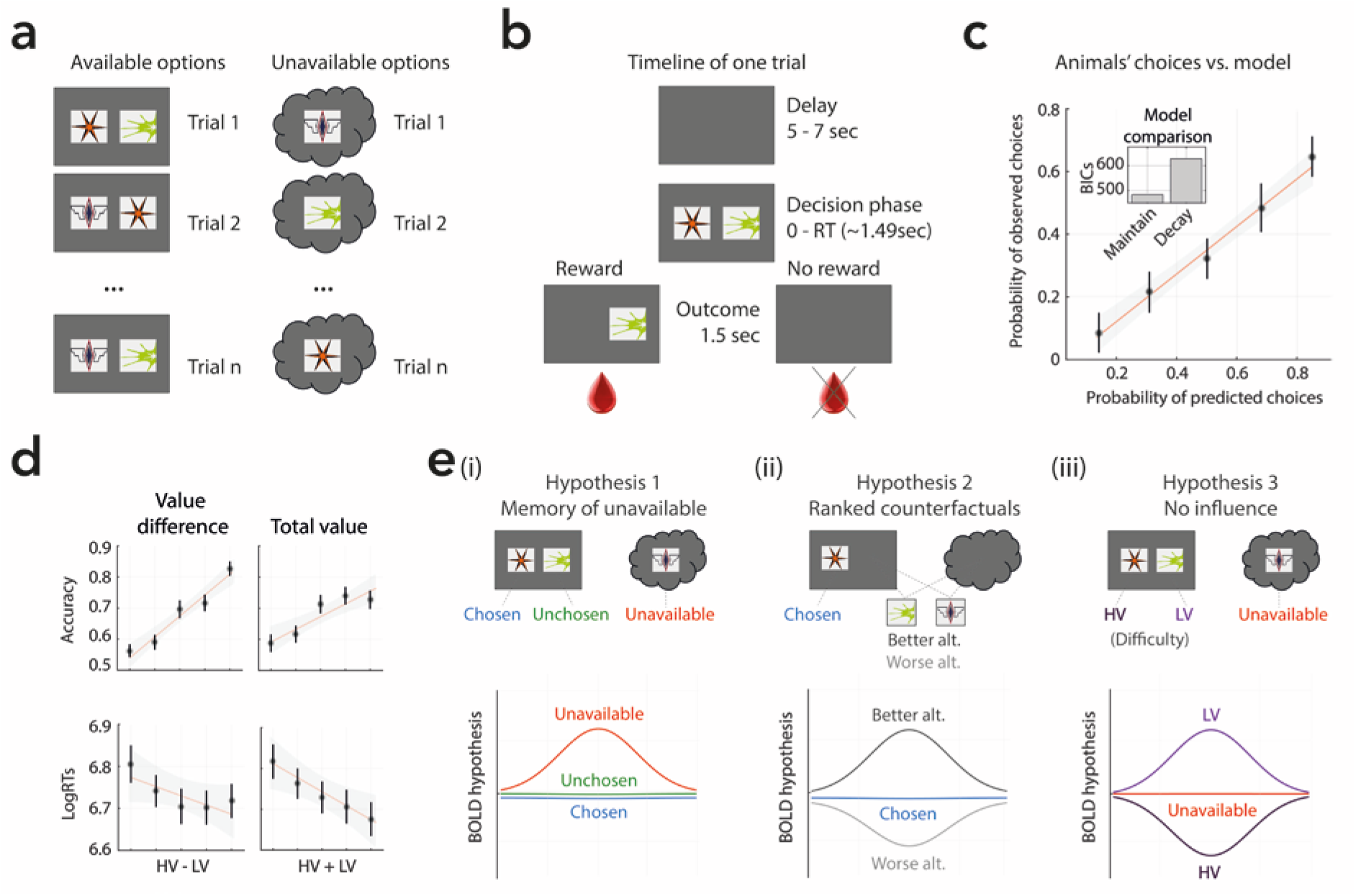
**(a)** On each trial, animals could choose between two symbols presented on the screen and had to keep in mind a third option, unavailable to them. The position of each symbol on the left/right part of the screen and the combination of available/unavailable options was fully and pseudorandomized respectively. **(b)** Each trial began with a random delay followed by the presentation of two abstract symbols for a period ending when the animals made a choice. During this time monkeys pressed one of two touch-sensors to indicate which of the two symbols (right or left) they believed was more likely to lead to a reward. Finally the decision outcome was revealed for 1.5 sec. The selected symbol was kept on the screen (or not) to inform the monkeys of a reward delivery (or no reward). **(c) Main panel:** Model-predicted choice probabilities (x-axis) derived from an RL algorithm (Maintain model) using the softmax procedure (binned into 5 bins – bin size of 0.2 – and averaged across all animals and across symbols) closely matched animals’ observed behavioral choices (y-axis), calculated for each bin as the fraction of trials in which they chose one of the three symbols. **Small panel:** A Bayesian model comparison using BIC scores revealed that the Maintain model explained the data better than a Decay model in which a free parameter captured how much the animals would “forget” the unavailable option. **(d)** The top graphs show the proportion of correct choices (selecting the option with the highest reward probability) plotted as a function of difficulty (distance between the better high value [HV] and the worse low value [LV] presented options: **left panel**) and context value (sum of both HV’s and LV’s expected values: **right panel**). Decision accuracy improved with higher value difference between available options and higher total value. The bottom graphs show log-transformed mean RTs (±SEM) plotted as a function of difficulty and context. LogRTs decreased for easier decisions and higher trial value. Solid lines are linear fits to the data and the shaded area is the 95% confidence interval. **(e)** Because each of the three options’ values were uncorrelated with one another it was possible to look for neural activity according to three main coding schemes. **[i]** If activity in a brain area covaries only with the value of the unavailable option then this suggests the area is concerned with representing the value of an option held in memory on the current trial and which should not interfere with decisions taken on the current trial. **[ii]** If instead activity covaries with the ranked value of both the unchosen available option and the option held in memory then it reflects the value of any currently counterfactual choice that might be taken in the future. It is important, however, to distinguish such a pattern from a third possibility **[iii]** in which neural activity is only reflecting the currently available options without representing the counterfactual or unavailable option. Thus the activity would be negatively related to the HV available option value and positively related to the LV option value. This third pattern indicates that the brain area’s activity reflects the difficulty or uncertainty of the current decision because the difficulty of selecting an option becomes harder as the LV option increases and as the HV option decreases but it is unaffected by the value of the choice that cannot currently be taken (see discussion by Kolling and colleagues^1,2^).

Behavioral analyses demonstrated that animals maintained representations of counterfactual choice values to guide future behavior on subsequent trials. We therefore used fMRI to test whether neural activity reflected counterfactual choice values according to one of several possible schemes. First, neural activity might simply represent the value of the unavailable option (Hypothesis 1). Alternatively, it might reflect the value of any counterfactual option – options that are currently unavailable for choosing, and options that are available on the current trial but which are unchosen. In such a scheme, it may not be important whether a counterfactual choice is unavailable or unchosen, however, if such a representation is to guide future behavior, then it should reflect the ranked values of the alternative options (Hypothesis 2). We also compared this with a third scheme in which an unavailable option’s value had no influence on neural activity (Hypothesis 3) (fig.1e). We found evidence for the existence of counterfactual values in ACC, anterior lateral prefrontal cortex (lPFC), and hippocampus. Activity in these regions was related to accurate decision making in future decisions but not in the current decision. The activity pattern in ACC further indicated that it was in this region that counterfactual values were transformed into actual behavioral change.

Because we used an animal model it was possible to investigate not just correlation between neural activity and behavior but the activity’s causal importance for behavior^3^. We examined the impact of transiently disrupting ACC neural activity with Transcranial Focused Ultrasound Stimulation (TUS). The TUS 250 kHz ultrasound stimulation was concentrated in cigar-shaped focal spot several centimeters below the focusing cone. Importantly, we recently demonstrated that this protocol transiently, reversibly, reproducibly, and focally alters neural activity (Verhagen et al., BioRXiv) and can do so even in a relatively deep area such as ACC (Folloni et al., BioRXiv). Additionally, a similar TUS protocol altered saccade planning in macaques when applied to the frontal eye fields but not to a location 10-12mm distant in dorsal premotor cortex^4^. In the current study, consistent with our ranked counterfactual hypothesis (Hypothesis 2), ACC TUS impaired translation of counterfactual choice values into actual behavioral change.

## Results

### Animals learned option values and maintained them in memory without forgetting

To behave adaptively in this task, animals should estimate each option’s reward probability and maintain these estimates in memory. If there are three options (for example A, B, and C) then animals should retain what they have learned about option C even if subsequent trials involved presentation of only options A and B. The representations of C’s reward value should then guide future decisions when C becomes available again. We therefore modeled animals’ choices using a reinforcement learner^4,5^ and tested whether the unavailable option’s estimated reward probability (which in our experiment determines expected value) was maintained in memory. Specifically, we used two reinforcement-learning models to simulate behavior: 1) the Maintain model which assumes the expected value associated with the unavailable option does not change or decay on trials when it is not shown^6–8^; 2) the Decay model in which memory of an unavailable option’s value decays or is “forgotten” as a function of time whenever it is not presented (Methods)^9,10^.

For both models, we estimated free parameters by likelihood maximization and Laplace approximation of model evidence to calculate the Bayesian Information Criterion (BIC) and the exceedance probability respectively (supplementary fig. 1; Methods). Bayesian model comparison revealed that monkeys did not forget unavailable option values (Lower BIC values indicate better model fit, Maintain model: BIC=486.80, Decay model: BIC=618.75; fig.1c). Exceedance probabilities for the models based on approximate posterior probabilities suggested the Maintain model significantly outperformed the Decay model (*ϕ*=0.96; see supplementary fig.1). Animals learned the options’ values and maintained them in memory without forgetting even when options were not available on a given trial.

To confirm the relationship between the better model’s predictions and behavior, we compared choice probabilities predicted by the Maintain model and the actual recorded frequencies of animals’ responses and found that the model matched behavior well (fig.1c). A further linear regression analysis at the individual level generated an average *R^2^* of 0.92 across animals and regression coefficients that were significant (P<0.005) for each animal. Having established the goodness of fit of the Maintain model to behavior, all further analyses were conducted using the expected values estimated with this model. To predict behavior as in humans and artificial decision making networks^11^, estimates for the two available options were categorized as “high value” (HV) and “low value” (LV) and accuracy was categorically defined as HV selection. With these estimates, we found that the difference in value between the two available options (sometimes called “difficulty” as depicted in fig.1e[iii]) as well as the total value of available options were reliable predictors of animals’ choice accuracy (value difference: t_24_=7.12, P<0.001; total value: t_24_=4.10, P<0.001) and reaction times (value difference: t_24_=-3.68, P=0.001; total value: t_24_=-5.54, P<0.001; fig.1d).

### Value associations of counterfactual options guide future choices

To guide future behavior, it is essential to retain counterfactual choice values in case these choices become available again in the near future. There are at least two different ways that animals can maintain counterfactual information for future use. The first way is to consider which choices are available and which are not on each trial (Hypothesis 1; fig.1e[i] left panel) and thus to categorize the options as “chosen”, “unchosen” and “unavailable”. A second way to describe the options (Hypothesis 2; fig.1e[ii] middle panel) is to think of both the unchosen and the unavailable options as alternative courses of action. Together, these two options constitute the counterfactual choices – potential choices that were not, or could not, be taken on the current trial but which might be taken in the future. Animals might rank the expected value associated with the counterfactual options. Therefore, instead of categorizing the options as “unchosen” and “unavailable”, we characterized them as the “better” and “worse” counterfactual options (also named “better” and “worse” alternative options), irrespective of their availability. Sometimes, the unchosen option will be the better alternative, and sometimes the unavailable option will be the better alternative. Finally we can test the hypothesis that animals do not maintain counterfactual information for future use but only represent information about counterfactual choices so as to represent the difficulty of the current decision (Hypothesis 3; fig.1e[iii] right panel). Consideration of this possibility is important because it has been suggested that activity in some of the brain areas that we investigated, such as ACC, reflects the difficulty of response selection^14,15^.

In line with the first hypothesis, we performed a logistic regression assessing whether the unavailable option’s expected value influenced its future selection when it next reappeared on the screen. This analysis revealed that decisions to select the previously unavailable option were strongly related to its expected value (t_24_=3.31, P=0.002; fig.2a), indicating monkeys held the memory of its value even over trials on which it was not available in order to guide later behavior, aligning with the prediction of the Maintain model. A complementary analysis confirmed these results and showed that accuracy of the future choice was influenced by the currently unavailable option, particularly when its most recent expected value was the best of the three options ⅛4=6.59, P<0.001; fig.2b) regardless of the last reward/choice association (paired t-test between rewarded/non-rewarded choices at t-1: P<0.001) and beyond the effect of the current HV and LV on the screen (all one-sample t-tests: P<0.001).

**Figure 2:**
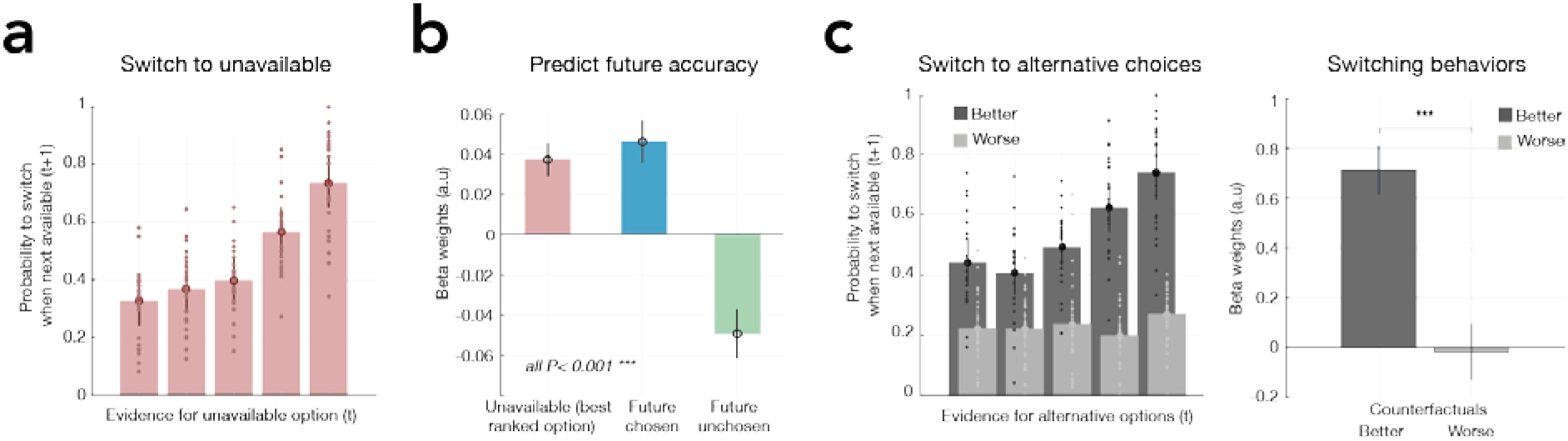
Future switches are explained by the expected value associated with counterfactual options. **(a)** Estimated expected values associated with the unavailable option on the current trial predict whether animals switch to it when it reappears on the screen on subsequent trials (y-axis: probability of switching to the currently unavailable option. x-axis: reward probability associated with the unavailable option estimated from the Maintain model). Each bin contains 20% of averaged data across sessions (±SEM) **(b)** A logistic regression confirms that accuracy is explained by the currently unavailable option’s value (higher accuracy, for trials in which it is the best of the three options vs. when it is not), in addition to the value of the future chosen and unchosen options (beta coefficients are averaged across sessions ±SEM) **(c)** A similar analysis to the one shown in panel **(a)** is performed but on the basis of a new coding scheme where the counterfactual options (current unchosen option and current unavailable option) are ranked according to their associated reward probabilities as the better and the worse counterfactual choices **(left panel)**. The value of the better counterfactual option significantly influenced the frequency with which monkeys subsequently switched to it but this was not the case for the worse counterfactual option **(right panel)**.

In line with the second hypothesis, we performed a series of analyses similar to those described above but replacing value estimates for the unavailable option by estimates for better and worse alternative choices. These analyses revealed animals’ decisions to switch to the better counterfactual choice were influenced by its expected value (t_24_=5.68, P<0.001) but this was not true for the worse counterfactual choice (t_24_=0.21), suggesting animals switch to the next best alternative more effectively than to the worse alternative (fig.2c, left and right panels). Overall, the results demonstrate two ways of categorizing the choices made in the task: either by classifying them as “available” and “unavailable”, or by considering the current chosen option in contrast to better and worse counterfactual choices. These frameworks guided our following analysis of fMRI data (fig.1e).

### Hippocampal activity predicts successful future choices when the unavailable option becomes available again

Having established that animals not only represent choice value information that cannot be used on the current trial, but exploit this information on pending trials, the first fMRI-related analysis explored the extent to which neural activity reflected this key feature: the expected value of the currently unavailable option (Hypothesis 1; fig.1e[i] left panel). We tested for voxels across the whole brain where activity correlated with the trial-by-trial estimates of the unavailable option’s expected value, particularly when the future selection was successful (GLM1: value of unavailable option on trial t when switching to it correctly on trial *t+1* vs. value of unavailable option on trial t when switching to it incorrectly on trial *t*+1). We also included the expected value of the chosen and unchosen options as separate terms in the GLM (Methods). This analysis revealed one region in which the neural value coding of the unavailable option was different for successful future selection compared with unsuccessful future selection, surviving multiple correction (Z>3.1, whole-brain cluster-based correction P<0.001): right hippocampus (peak Z=3.61, Caret-F99 Atlas (F99): x=16.5, y=- 7.5, z=-12). At a lower threshold, we also found its bilateral counterpart: left hippocampus (peak Z=3.05, F99 x=-14, y=-9, z=-12.5; fig.3a). To illustrate the significant activity in bilateral hippocampal regions, we extracted the time course of the neural activation in two regions of interest (ROIs), defined as spheres centered on the peak of the activations (fig.3b left). Note that this analysis was performed for illustrative purposes only as the ROIs were formally linked to the comparison between correct and incorrect future selection used to establish the ROI location^16^. The activity pattern represented in this analysis is noteworthy as it shows that the blood oxygenation level dependent (BOLD) signal in hippocampus is scaled by the expected value associated with the unavailable options only when the currently unavailable option is going to be chosen correctly on a future trial (see supplementary fig.2).

**Figure 3.**
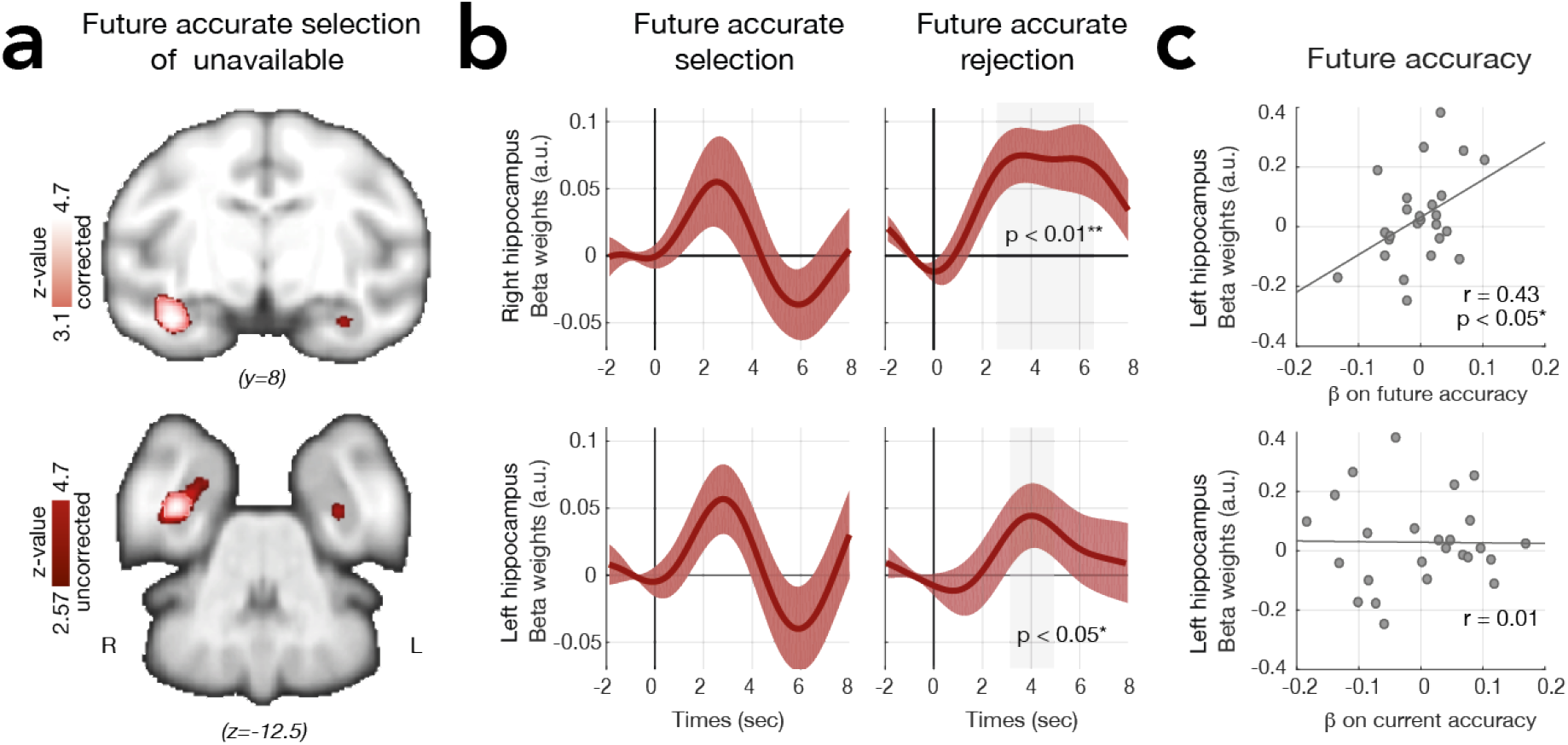
Unavailable option value signal in hippocampus favors accurate future planning. **(a)** A whole-brain analysis tested for voxels where activity correlated with the trial-by-trial estimates of the unavailable option binned according to successful future selection. The fMRI analysis was time-locked to the decision phase on trial t and binned according to accurate vs. inaccurate selection of the unavailable option on trial t+1 (in light pink: cluster-corrected, Z > 3.1, P < 0.001; in red: uncorrected) **(b)** ROI analyses of the right (**top panels**) and left (**bottom panels**) hippocampus illustrate the time course of the aforementioned contrast. BOLD fluctuations reflect the value of the unavailable option on the current trial when it is accurately versus inaccurately selected on the next trial (**left panels** illustrate the contrast show in **(a)**). A leave-one-out procedure (for spatial and temporal peak selection) to assess statistical significance revealed that a similar activity change occurs when contrasting the value of the unavailable option for accurate versus inaccurate future *rejections* of the unavailable option (right panels). **(c)** In the left hippocampus, the beta weights for the contrast used in **(a)** and illustrated in **(b, left panel)** were predictive of how much the unavailable option’s reward probability influenced animals’ future choice accuracy **(top panel)** but this was not true for current choice accuracy **(bottom panel)**. Results are normalised.

The hippocampus’ role in maintaining information about currently unavailable choices for prospective guidance of future decisions might not be restricted to occasions when that choice will be made in the future; it may also encompass the prospect of rejecting the currently unavailable option if it is likely to be worse than the others^17^. To demonstrate this, we repeated the analysis in the trials preceding those in which the animal decided *not* to select a currently unavailable option. Critically, this analysis also revealed a greater BOLD signal for the value of the unavailable option on the current trial when this option was correctly rejected in the future compared to when it was incorrectly rejected (Leave-one-out peak selection: right Hippocampus: P=0.006, t_24_=2.96; left Hippocampus: P=0.03, t_24_=2.19; fig.3b right). In summary, hippocampal activity is scaled by the currently unavailable option’s value more strongly (e.g. there is a stronger memory trace) when the next decision involving that option is going to be made correctly regardless of whether it is going to be chosen correctly (because it is highest in value) or rejected correctly (because it is lowest in value) in the future.

Finally, having established that hippocampal activity is related to memory of unavailable options we hypothesized that variation in such activity (at trial t) across sessions might predict variation in influence of the unavailable option’s value on future accurate switching behavior (at *t+1*) (fig.2b). We found a significant correlation in the case of future decisions in which the unavailable option became accessible (r=0.43, P<0.05) but there was no correlation for the current decision while the unavailable option remained inaccessible (r=0.01, P>0.5; fig.3c). This result again suggests that the hippocampus is involved in future planning but not current, on-going decision making, as further confirmed by connectivity analysis with regions involved in guiding the ongoing decision (see supplementary PPI analysis).

### ACC ranks counterfactual options according to their expected value

The previous analysis was predicated on the idea that the brain maintains information in memory pertaining to currently unavailable choices while encoding what is relevant for the current decision (the two available options) elsewhere in the brain. Therefore, we next sought brain regions encoding the key decision variable – how much better is the currently chosen available option compared to the currently rejected available option. We searched for activity that parametrically encoded the difference in value between the currently chosen and unchosen options (GLM2: subjective choice comparison model with chosen vs. unchosen expected values). Such a neural pattern, when locked to decision time, is sometimes referred to as a choice or value comparison signal. We found strong bilateral activations in a distributed network including ACC (peak Z=-3.75, F99 x=1, y=20.5, z=10.5), lPFC (right peak Z=-4.61, F99 x=14.5, y=17.5, z=9.5; left peak Z=-4.29, F99 x=-15, y=16, z=9.5) and ventromedial prefrontal cortex and adjacent medial orbitofrontal cortex (vmPFC/mOFC; peak Z=-4.01, F99 x=-5, y=14, z=2) encoding the (negative) difference in expected value between the chosen and unchosen options (fig.4a; |Z|>3.1, whole brain cluster-based correction P<0.001). In other words, activity in these areas increased as decisions became harder (e.g., because the subjective value of the chosen option became lower or the subjective value of the unchosen option became higher or both).

**Figure 4.**
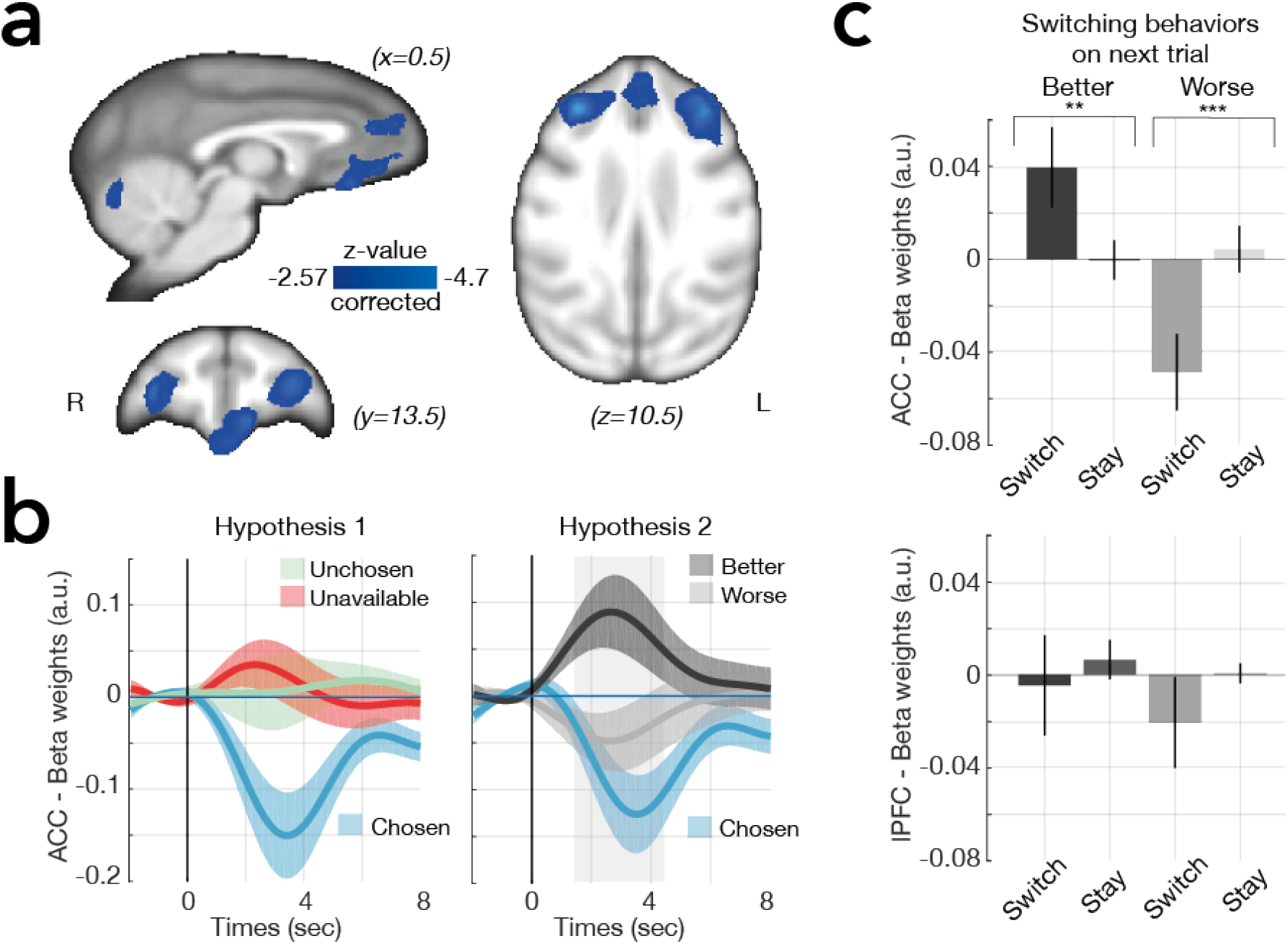
The anterior cingulate ranks expected reward probabilities of counterfactual options. **(a)** Whole-brain analysis shows a significant negative relationship between BOLD activity and the difference between the expected value associated with the currently chosen and unchosen options in a distributed brain network, including ACC, bilateral lPFC, and vmPFC/mOFC (cluster corrected, Z > 3.1, P < 0.001) **(b)** ROI analysis of the ACC illustrates the relationship between BOLD and the fully parametric representation of the currently chosen, unchosen, and unavailable options (left panel) and shows that a distinct model in which the counterfactual options are ranked according to their associated reward probabilities explains the data better. Note that we avoid double dipping in favor of the hypothesis that we want to support (hypothesis 2) since the ROI has been defined on the basis of hypothesis 1. **(c)** The parametric representation of the better and worse counterfactual values in ACC was further explained by whether a future switch in behavior will occur as opposed to the continued maintenance of behavior (“stay”) (leave-one-out procedures for peak selection on time series analyses: **top panel**). This was not true in the lPFC (**bottom panel**)

To first illustrate the relationship between option values and lPFC and ACC activity, we extracted BOLD time courses (using a leave-one-out cross-validation approach to avoid circularity of analyses) from ROIs over each region and performed further analyses (Methods). For each region, we found activity related to the key decision variable – the difference between chosen and unchosen values – was mainly driven by the negative relationship of the BOLD signal with the expected value of the chosen option (all |Z|>3.1 for the Chosen regressor); there was no effect related to the unchosen option value (no significant activity for the Unchosen regressor). Importantly, the analysis contained an extra regressor representing the unavailable option’s value, which also had no significant effect (no significant activity for the Unavailable regressor), suggesting that, aside from the choices that the animals had experienced directly, the representation of the other two options as “unchosen” and “unavailable” provided a poor account of activity in these regions (ACC and lPFC). Importantly, the negative relationship between the ACC BOLD signal and the value of the chosen option may reflect the opportunity cost of switching away from the current choice.

Following this idea, in a second step, we tested whether the ACC might represent the possible alternatives that the animal might switch to in the future (Hypothesis 2 in fig.1e middle panel). In this scheme, the two options not selected on the current trial, the unchosen option and the unavailable option, could both be considered counterfactual options that might be taken in the future and which could be ranked according to their expected value (GLM3: better vs. worse alternatives model, as per our behavioral analyses). Using Bayesian statistics for each region within the same network (see Methods), we found that the activity pattern representing better and worse alternatives provided a significantly better account of neural activity in both ACC and lPFC compared to either the subjective choice comparison model (GLM2) or a third model (GLM4) that does not represent alternative options but rather the difficulty of selecting the current response (Hypothesis 3 in fig.1e) with *ϕs*>0.95 (fig.4b and see supplement fig.3 and methods for full Bayesian Model Comparison^18^). Thus, ACC and lPFC activity parametrically scales with the values of counterfactual choices. One interpretation of the activity pattern is that it forecasts choosing the better of the counterfactual options during future decisions.

We directly tested this hypothesis using multiple regressions to investigate whether the activity in lPFC or ACC would predict upcoming switching behavior. For each ROI, we employed four regressors time-locked to the stimulus period of trial t, including i) the expected value of the better alternative if the future trial is a switch to that option; ii) the expected value of the better alternative if the future trial is a stay (i.e. repetition of the same choice as on the current trial); iii) the expected value of the worse alternative if the future trial is a switch to that option; iv) the expected value of the worse alternative if the future trial is a stay. ACC activity predicted upcoming decisions to switch to the better and avoid the worse counterfactual (fig.4c; leave-one-out procedures for peak selection: ANOVA: main effect of area: F24=6.15, P=0.02; *post-hoc tests:* Best: t_24_=2.41; P=0.02, Worst: t_24_=-2.94, P=0.007) but this was not true in lPFC (all Ps>0.2). Such a pattern is consistent with a role for ACC in evaluating future strategies before execution^8,19–21^. By contrast, macaque anterior lPFC holds estimates of counterfactual choice values that are less immediately linked to behavior. Similarly, human frontal polar cortex activity reflects the values of alternative choice strategies in a manner that is also less immediately linked to behavior^22^.

There has been considerable interest in the possibility that ACC activity simply reflects decision difficulty, which is inversely proportional to the difference in value between the option chosen and the available option left unchosen^4^ (fig.1e, right panel). When one option’s value is much higher than the other option, the decision is easy. But when the values of the two options are similar, the decision is difficult because it is hard to reject an alternative that is close in value. However, ACC activity cannot be explained by difficulty because GLM2 demonstrated it did not reliably reflect the value of the rejected available choice and GLM3 demonstrated that instead it reflected the best counterfactual choice value even when that choice was not currently available.

### ACC disruption impairs translation of counterfactual choice values into actual behavioral change

To test whether counterfactual choice value representations in ACC were causally important for effective behavioral switching, TUS was applied to the same ACC region in two of the four macaques. We previously demonstrated, using resting state fMRI data that 40s sonification at 250 kHz reaches ACC and does so in a relatively focal manner having less effect on adjacent, even overlying, brain areas (Folloni et al., BioRXiv). The same stimulation was applied to ACC using MRI-guided frameless stereotaxy^22,23^ immediately prior to nine testing sessions that were interleaved, across days, with nine control sessions in which no TUS was applied in each monkey (fig.5a; Methods). As in all previous fMRI sessions, novel stimulus images were assigned to options at the start of each day’s testing session although the underlying reward environment remained constant. While there were clear differences in choice patterns between the ACC TUS and control conditions (fig.5b) these did not result in straightforward accuracy differences (fig.5c; P > 0.5).

**Figure 5.**
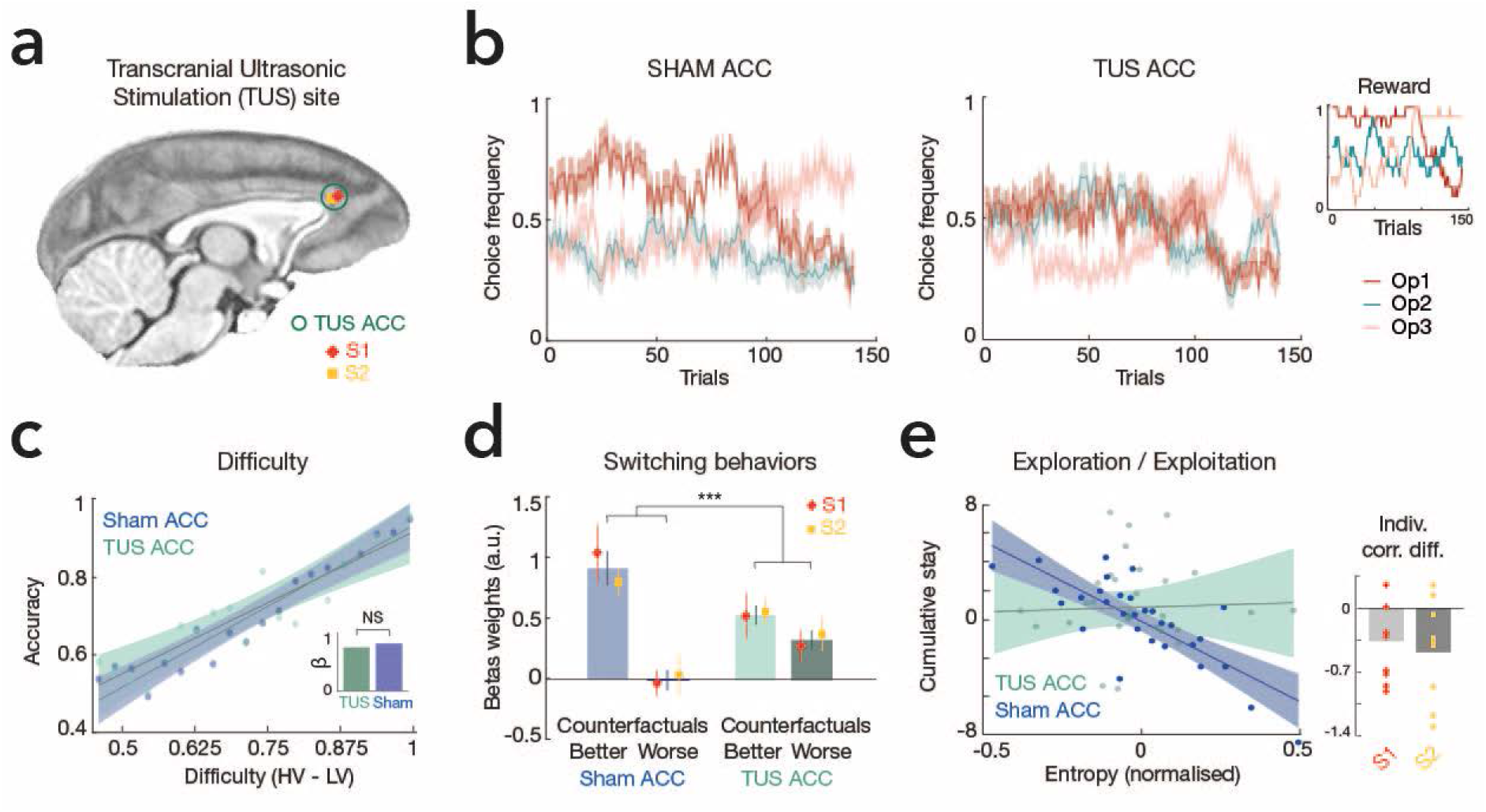
Transcranial Focused Ultrasound Stimulation (TUS) of ACC impaired translation of counterfactual choice values into actual behavioral change. **(a)** Neurostimulation site where TUS was applied for each animal (S1 and S2). The TUS transducer was set at a resonance frequency of 250 kHz and concentrated ultrasound in cigar-shaped focal spot in ACC. **(b)** Running average choice frequency for the three options in the control/sham ACC **(left)** and the TUS ACC condition **(middle)** across sessions. Predetermined reward schedules used in the sham and in the TUS ACC task for three options, similar to the task used in the fMRI experiment **(right)**. **(c)** Decision accuracy (selecting the option with the highest reward probability) plotted as a function of difficulty (difference in value between the best [HV] and worst [LV] presented options). There was no difference between the TUS ACC and the control conditions. Solid lines are linear fits to the data and the shaded area is the 95% confidence interval. **(d)** The significant difference between the influence of the better and worse counterfactual option value on future switching behavior (in blue, as per fig.2f) was significantly reduced after TUS ACC (in green). **(e)** While entropy (summed entropy of reward probability for all options) is strongly and negatively predictive of a change in exploratory behavior in the control condition (indexed by the cumulative number of “stay” choices: choices of the same option on one trial after another), this relationship is disrupted in the TUS ACC condition. The small panel depicts the difference between the TUS ACC and the sham condition for each pair of days (Animals 1 [S1] and 2 [S2] are individually represented as red diamond and yellow square, respectively in all plots).

To test formally whether ACC translates counterfactual choice values into actual behavioral change, as in the earlier behavioral analyses (fig.2f), we regressed the frequency with which monkeys switched, on one trial, onto the values of choices that, on a previous trial, had been counterfactual alternatives (fig.5d). As in previous analyses, without TUS, the value of the better counterfactual option significantly influenced the frequency with which monkeys subsequently switched to it (P<0.001; t17=6.5) but this was not the case for the worse counterfactual option (P=0.9; t_17_=-0.06). This was, however, not true for the TUS condition. When comparing the control with the TUS data, a twoway ANOVA revealed a significant interaction between the effect of TUS and the influence of counterfactual values on switching behavior (P=0.002; F_17_=12.97). The significant difference between the influence of the better and worse counterfactual option value on future switching behavior that was present in the baseline condition (post hoc test: P<0.001; t_17_=7.56) was abolished (P=0.11; t_17_=1.7) after ACC TUS.

We further hypothesized that this behavioral change would impact the monkeys’ search strategies^13^ and reduce the influence of entropy (the unpredictability of the environment; see Methods for the computational definition of entropy) on their exploratory behavior^24^. In a running window analysis, we used the slope of entropy to predict the slope of cumulative stay choices (i.e. successive choices of the same option)^25^. As lower entropy favors exploiting knowledge to maximize gains and higher entropy favors exploring new options and discovering new outcomes, we expect to see a negative relationship between entropy and the frequency of stay choices. In the control group condition, we found such a relationship (P<0.001; t28=-6.59) but this was not the case after ACC TUS (P=0.82; t_28_=0.22).

In a final TUS experiment, to control for the anatomical specificity of the observed effects, we examined the effect of TUS to lateral orbitofrontal cortex (lOFC), a brain region also associated with distinct aspects of reward-guided learning and decision making^26,27^. LOFC TUS, however, had no impact on the way in which counterfactual choice value was translated into subsequent actual behavioral switching (supplementary fig.4).

### The unavailable option value affects the current value comparison via vmPFC/mOFC

So far, we have reported evidence that activity in a distributed circuit spanning hippocampus, anterior lPFC, and ACC reflects counterfactual choice values. Hippocampal activity reflects values of currently unavailable options while lPFC and ACC activity reflects the value of the current, better counterfactual choice regardless of availability. In ACC, counterfactual choice values are translated into behavioral change and disruption of ACC activity impairs the translation process.

One other area, vmPFC/mOFC, also carried a choice value comparison signal (fig.4a and fig.6b). This pattern of decision-related fMRI activity in vmPFC/mOFC has been reported previously in macaques^27^. Not only does BOLD activity increase as the decision becomes harder but single neuron activity also appears to reflect a decision process^26–29^ and vmPFC/mOFC lesions disrupt value-guided decision making^30^. The activity is in a location^27,31,32^ corresponding with that in which similar signals, albeit with an inverted sign, encode values of chosen and unchosen options during decision making in humans^13,33–36^. Given vmPFC/mOFC’s importance for many aspects of decision making^27,30^, it is noteworthy that unlike ACC, vmPFC/mOFC activity reflecting better and worse counterfactual values did not predict behavioral switches on future trials (as per results presented in fig4c, P>0.05). Instead, vmPFC/mOFC is concerned with the decision being taken now rather than decisions that might be taken in the future. In the following analyses, we tested whether the value of the unavailable option, rather than being maintained and leading to future successful switches, modulated the current decision between available options via vmPFC/mOFC.

**Figure 6.**
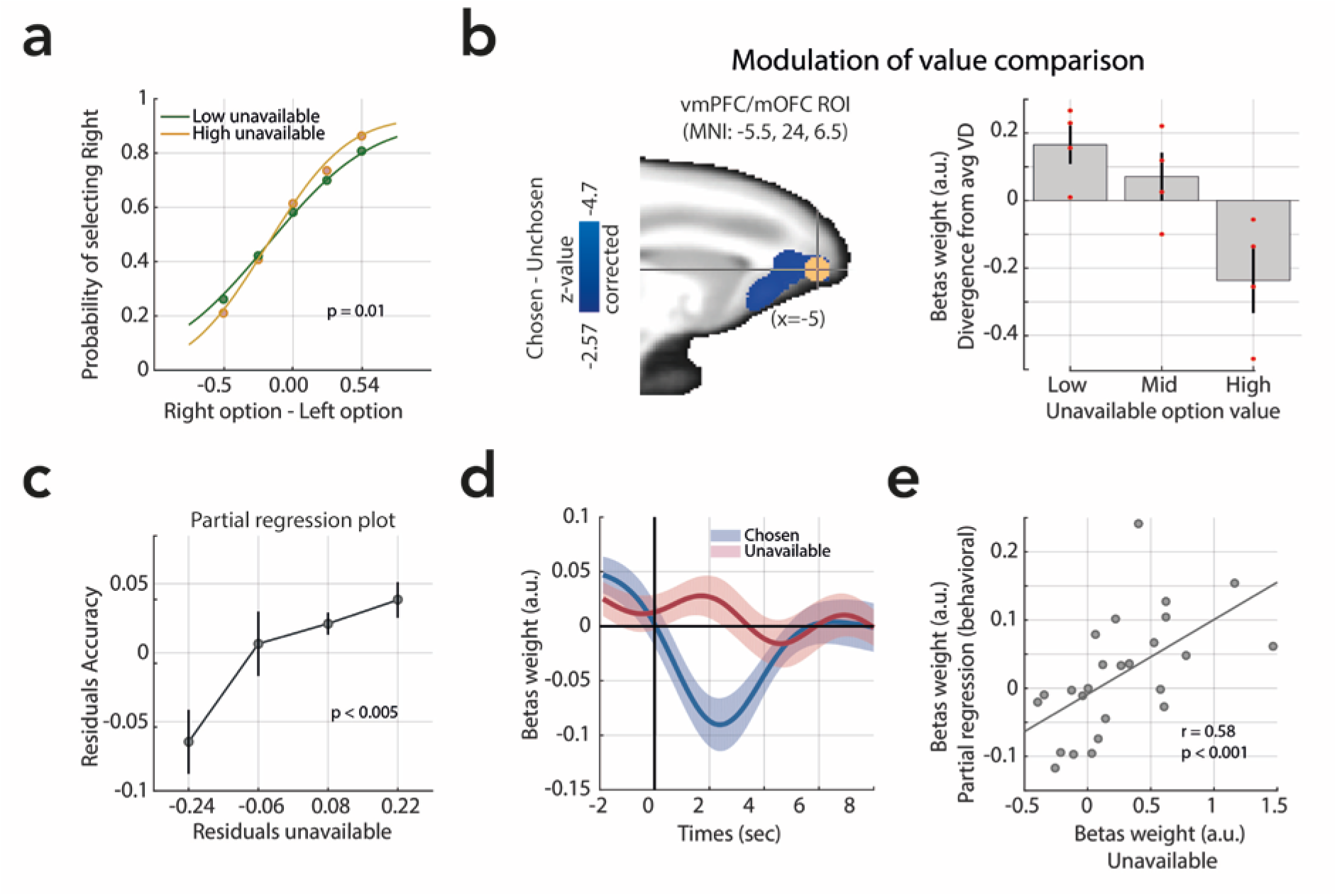
Contextual modulation of value-guided choice. **(a)** Average choice behavior when choosing between the Left and Right options plotted as a function of the value of the unavailable option (low: green; high: yellow). Decisions were less accurate when they were made in the context of a low value unavailable option. Curves plot logistic functions fit to the choice data. **(b)** ROI analysis of the vmPFC/mOFC (left panel: ROI sphere) illustrates the relationship between the BOLD value-comparison signal and the expected value associated with the unavailable option (binned in Low/Mid/High) (right panel). The greater the value of the unavailable option, the more negative the value difference; a more negative pattern is normally associated with decisions that are easier to take (see panel d). Data for individual animals are indicated by red dots. **(c)** A partial regression plot shows the uncontaminated effect of the unavailable option’s value on accuracy (y-axis: accuracy residuals; x-axis: residuals of the unavailable option’s value). Each bin contains 20% of averaged data across sessions (±SEM). (d) ROI time course analysis of the vmPFC/mOFC illustrates the relationship between BOLD and the fully parametric representation of the currently chosen and unavailable options **(e)** While there was not a main effect of the unavailable option value, vmPFC/mOFC variation in activity related to the currently unavailable option’s value explains between-session variation in the currently unavailable option’s influence on decision making. Scatter plot at the time of the peak effect of unavailable option value in the vmPFC/mOFC (leave-one-out peak selection).

We first assessed whether the unavailable option’s value influenced monkeys’ choices between available options. We computed accuracy (HV selection) and used a logistic regression to predict this categorical variable as a function of the unavailable option’s value (including HV and LV in the model). Our results strikingly show that the higher the value of the unavailable option, the better animals were at discriminating between the two available options (t_24_=3.79; P<0.001; similar results were obtain using a mixed-effect logistic regression model including sessions and animals as random effects using the *lmer4* package in the R environment: *χ*^2^_*(1)*_=25.78; P<0.001). To illustrate this effect, we represented frequency of choosing an option (for example the Right option) as a function of the value difference between the two available options (Right-Left option values) for two different levels of the unavailable option values (high vs. low; median split). Importantly, although the unavailable option can never be chosen, its value profoundly affects the efficiency of choice behavior. In all monkeys (see supplementary fig.5; test on *softmax* parameters; all Ps<0.05) and the group (fig.6a; t_24_=-2.66, P=0.01), relative choice curves were steeper when the unavailable option had high versus low values. In summary, unavailable options affected decision making; low value unavailable options rather than high value unavailable options were associated with lower accuracy. The result resembles a report that decision making between two options is improved if a third option, that is visually presented and cannot be chosen, is high rather than low in value^35^. In the previous study, the influence of the third option on decision making was mediated by vmPFC/mOFC. It is possible that a high value third option increases the total sum of inhibitory activity and that inhibitory activity drives a competitive selection process between the other two options^35^. Therefore, higher unavailable option value should result in a stronger value comparison signal in the vmPFC/mOFC associated with greater behavioral accuracy. We tested whether the same was true in the current data.

To examine vmPFC/mOFC activity independently from biases in peak selection, we used a literature-based ROI selection (in area 11m/11; fig.5b, left). We focused on activity reflecting the value difference guiding decisions between available options (chosen value–unchosen value) and binned it according to the value of the unavailable option (low: 0-33%; middle: 33-66%; high: 66-100% percentiles of unavailable option value). The vmPFC/mOFC response to the chosen value–unchosen value difference was modulated by the currently unavailable option’s value (linear mixed-effect analysis: t_10_=-4.01, P=0.002; fig.6b, right panel), in exactly the same way as behavior. Normally vmPFC/mOFC activity reflects the value of the chosen option with a negative sign (fig.4b and fig.6d); as the chosen option’s value falls and choosing it becomes more difficult, there is more activity in vmPFC/mOFC. This negative signal was diminished when the unavailable option value was very low and decisions between available options were less accurate. In summary, low (high) value unavailable options weakened (strengthened) the vmPFC/mOFC value comparison signal and weakened (strengthened) current decision accuracy. Importantly, the same analysis in the ACC and lPFC (both hemispheres) shows that the other areas behave differently and did not represent such modulation of value comparison by the unavailable option (all Ps>0.25).

To further test the strength of the link between the unavailable option’s impact on the current decision and its neural impact in vmPFC/mOFC we exploited variability in the behavioral effect across sessions. We hypothesized that variation across sessions in the size of the influence of the unavailable option’s value on vmPFC/mOFC would be related to variation in behavioral accuracy. To test this hypothesis, we first performed a partial regression analysis to reveal the uncontaminated effect of the unavailable option’s value on accuracy after controlling for the effects of the available options’ values (t_24_=2.84, P=0.008; see fig.6c). Separately, we extracted the unavailable option’s value-related signal change across sessions (time course analysis performed with the GLM2, see fig.6d for illustration of the chosen and unavailable options). Sessions with a greater unavailable option value-related signal in the vmPFC/mOFC also exhibited a higher effect of the unavailable option on accuracy in the current trial (r=0.58, P<0.001, see fig.6e).

## Discussion

Decision making is not only guided by accumulation of sensory evidence in favor of one choice over another but also by the values associated with choices that are currently unavailable but stored in memory^2^. It is both essential and a burden to store counterfactual choice values when other choices are actually being taken at the current point in time. On the one hand, it is essential to retain counterfactual choice values to *guide future behavior;* choices that are currently counterfactual may be taken in the future if they become available again, if the value of the choice currently taken diminishes, if the current choice is no longer available, or if the value of the counterfactual choice exceeds that of other alternatives offered in the future. We found that the four monkeys indeed held counterfactual choice values and when they used them to guide their decision making they were able to switch more effectively between choices and harvested more reward (fig.2b). On the other hand, holding information about counterfactual choice values is a burden because it *distracts from the current choice* to be taken; the monkeys chose less effectively between two available options when they had recent experiences of a currently unavailable low value option (fig.6a). One way of interpreting this finding is that both available options are much better than the unavailable option with which they are compared and so the monkeys choose either available option. Ecologically, this might be adaptive as an animal can afford to slow down in a high value environment, rather than having to satisfice quickly. As well, low (high) value unavailable options may reduce (increase) the net amount of inhibitory activity necessary for mediating the competitive selection process between available options and this leads to less/more accurate decision making. The benefits and costs of holding counterfactual choice values in memory were, respectively, associated with a neural circuit spanning hippocampus, ACC, and anterior lPFC (fig.3, 4, 5) and, on the other hand, with vmPFC/mOFC (fig.6).

The hippocampus-ACC-lPFC circuit suggested by the results is consistent with the known connections of these areas. The hippocampal formation interconnects with an identical ACC region via a fornical pathway from the subiculum^37^ while it interconnects with adjacent parts of lPFC via an extreme capsule pathway from presubiculum and parahippocampal cortex^38^. ACC and lPFC are strongly interconnected^39^ and frequently co-active^40–43^. The ACC is a key node in this network. It translates counterfactual choice values into actual changes in behavior; its coding of counterfactual choice values predicted whether monkeys would switch to those choices on a future trial (fig.4) and ACC disruption impaired the guidance of behavioral switching by counterfactual choice values (fig.5).

There has recently been debate about the degree to which ACC activity is simply explained by difficulty of response selection; ACC activity increases as decision difficulty increases^15,44^. However, in macaques, fMRI-recorded activity in other brain areas such as vmPFC/mOFC has the same property, it increases with decision difficulty. Nevertheless, macaque vmPFC/mOFC encodes the same information about choices as human vmPFC/mOFC even though human vmPFC/mOFC activity decreases with difficulty^27^ (figs.3, 6). It is therefore unlikely to be correct to attribute a role in difficulty detection to any area with activity that increases with decision difficulty. More direct evidence for this comes from the present study where current decisions improved as the unavailable choice value increased and ACC activity increased too (fig.4). Instead, the pattern of results is more in line with a view of ACC emphasizing the encoding of the value of alternatives^8,14,45,46^ and in mediating exploration of possible alternative choices^41,47,48^ as contexts change^42^.

Like the ACC, lPFC held counterfactual choice values but its activity was not closely related to behavioral change on future trials. In this respect, lPFC activity resembles that seen in parts of the frontal pole in humans that is also less directly related to behavior^14,33^. The lPFC region studied here may not, however, be homologous with the human frontal polar cortex but it may share some of the connections that the human lateral frontal pole has with other brain regions^31^. Holding counterfactual choice values in mind while other decisions are taken also compromises the accuracy with which the current decision is taken. Although the presence of high value distracting information can impair decision making via a process of divisive normalization of choice values,^35,49^ so can distracting low value choice information^11,35^. The two effects may depend on the distinct manner in which choices are encoded in intraparietal cortex and vmPFC/mOFC and damage to one area increases the distracting influence mediated by the remaining area^30,50^. In the present study, low value unavailable options were associated with diminished vmPFC/mOFC representation of the value difference relevant for guiding the current decision. Thus, even memories of choice values, as opposed to visually presented choice options, can impact on the accuracy of decision making via vmPFC/mOFC.

---

**Supplementary Figure 1:**
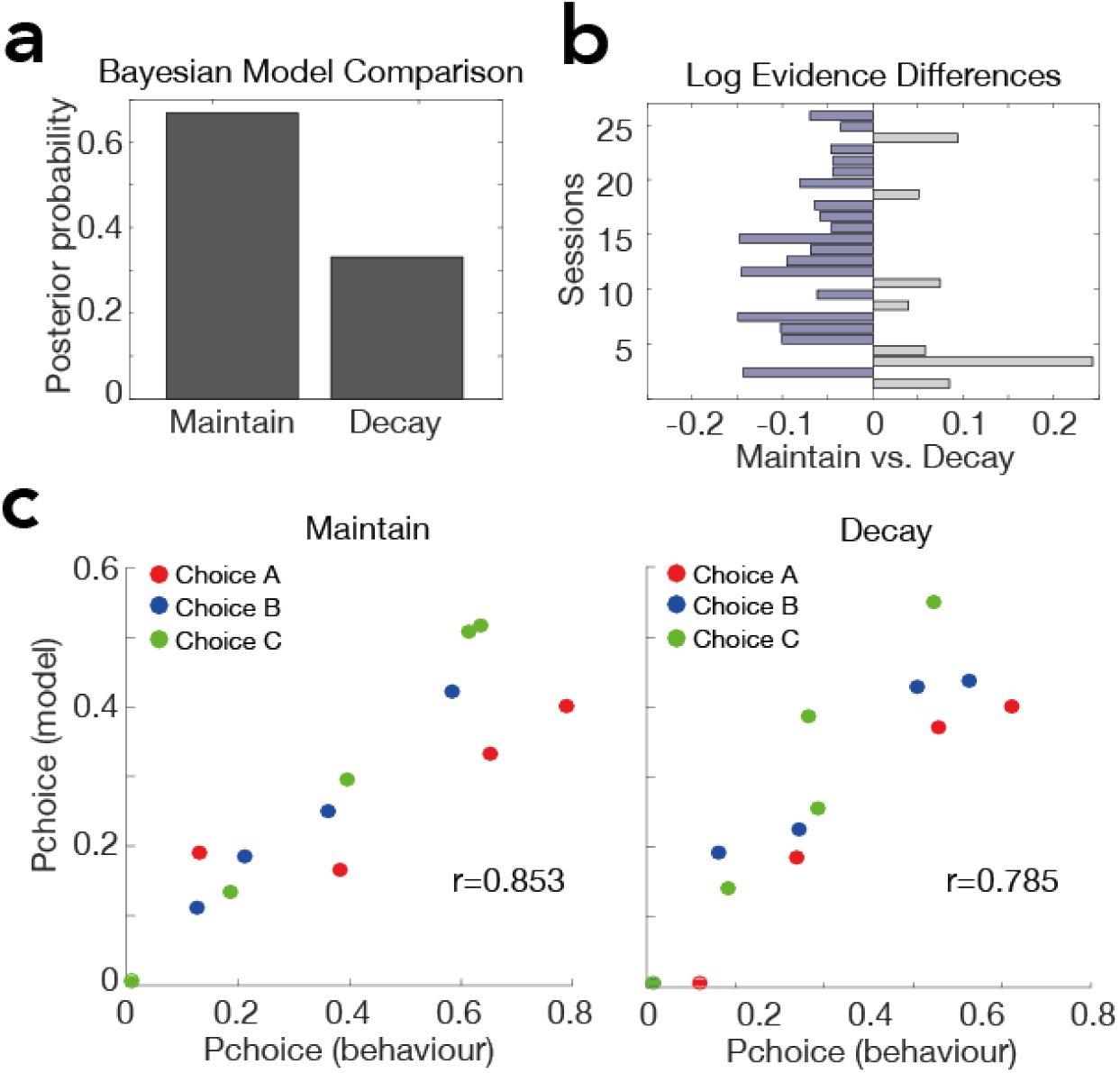
Model fit and model comparison. **(a)** To explain behavioral data, two models were fitted and compared: (i) a classical reinforcement learning algorithm that assumes no decay for the unavailable option (Maintain) and (ii) a modified version that assumes a decay for the unavailable option (Decay). Bayesian statistics revealed that there was a higher posterior probability that the Maintain model had generated the data than the Decay model. **(b)** The difference in log-evidences for all twenty-five data sets is plotted as a bar chart. **(c) Left and right panel:** Model-predicted choice probabilities (x-axis) derived from the Maintain and the Decay models respectively (binned into 5 bins – bin size of 0.2 – and averaged across all animals for each symbol separately).

**Supplementary Figure 2:**
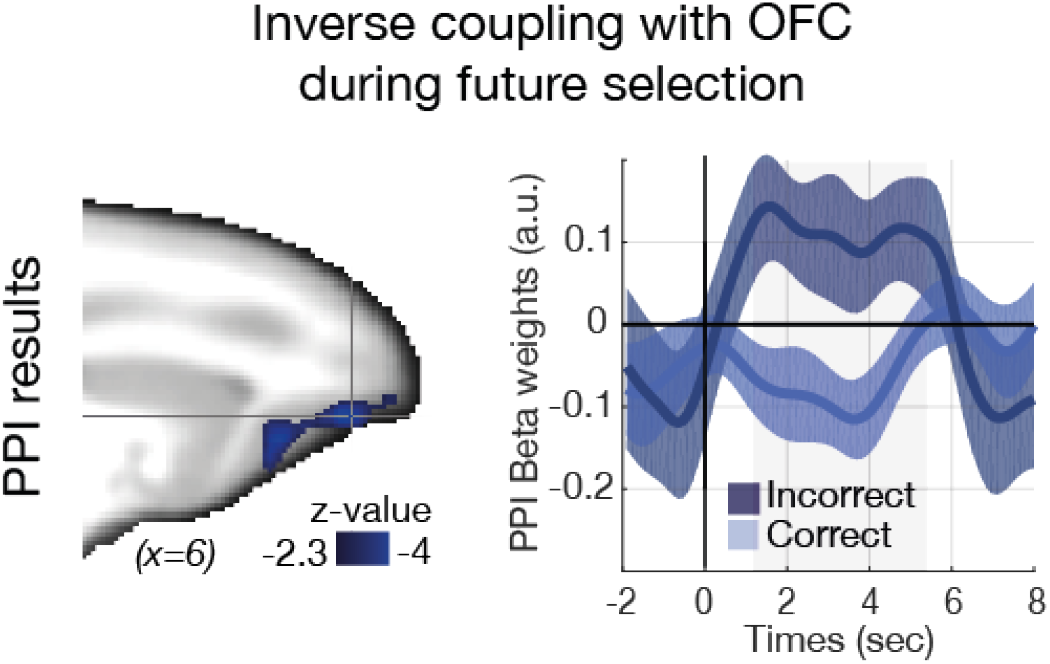
A hippocampal-frontal circuit in macaques holds memories of currently unavailable choice values to guide future behavior. **(a)** PPI results show that on the subsequent trial, when the previously unavailable option is correctly selected versus when it is incorrectly selected, the vmPFC/mOFC and right hippocampus are negatively coupled as a function of the currently unavailable option’s value. A negative coupling pattern is consistent with evidence that macaque mOFC/vmPFC codes the value of the choice that is taken with a negative change in activity (this means that it is more active when the decision is difficult because the choice taken is low in value).

**Supplementary Figure 3:**
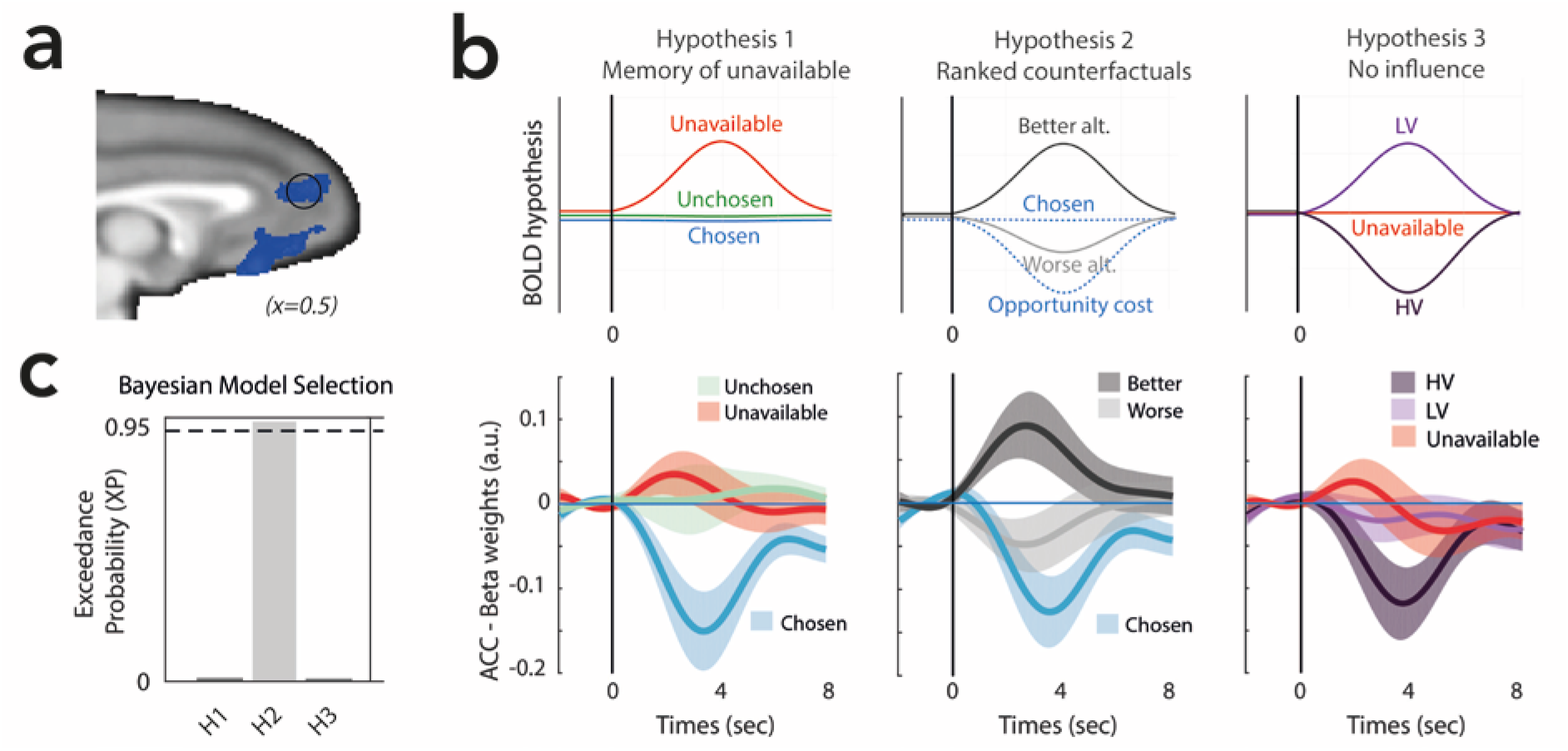
ROI analysis of the ACC illustrates the relationship between BOLD and the fully parametric representation of the estimates from the learning model that could be categorized in relation to the three hypotheses for action value coding scheme (see also fig.1e). **(a)** Time series were extracted from an ROI sphere centered on the ACC with a leave-one-out procedure. In the background is the result of the whole-brain analysis presented in the main manuscript in figure 4a. **(b)** The first coding scheme represents the currently chosen, unchosen, and unavailable options (left panel). The second coding scheme, which represents the counterfactual options ranked according to their associated reward probabilities, explains the data better (middle panel). The last model represents the HV option, LV option (available to the animal) as well as the unavailable option (right panel). **(c)** A formal model comparison showed that the 2^nd^ coding scheme outperformed the other two models. Bayesian model comparison of the three GLMs presented in the hypotheses section was performed within the functional ROI. Note that ROI selection avoids double dipping in favour of the hypothesis we aimed to validate, since the ROIs were defined from Hypothesis 1 that we aimed to reject.

**Supplementary Figure 4.**
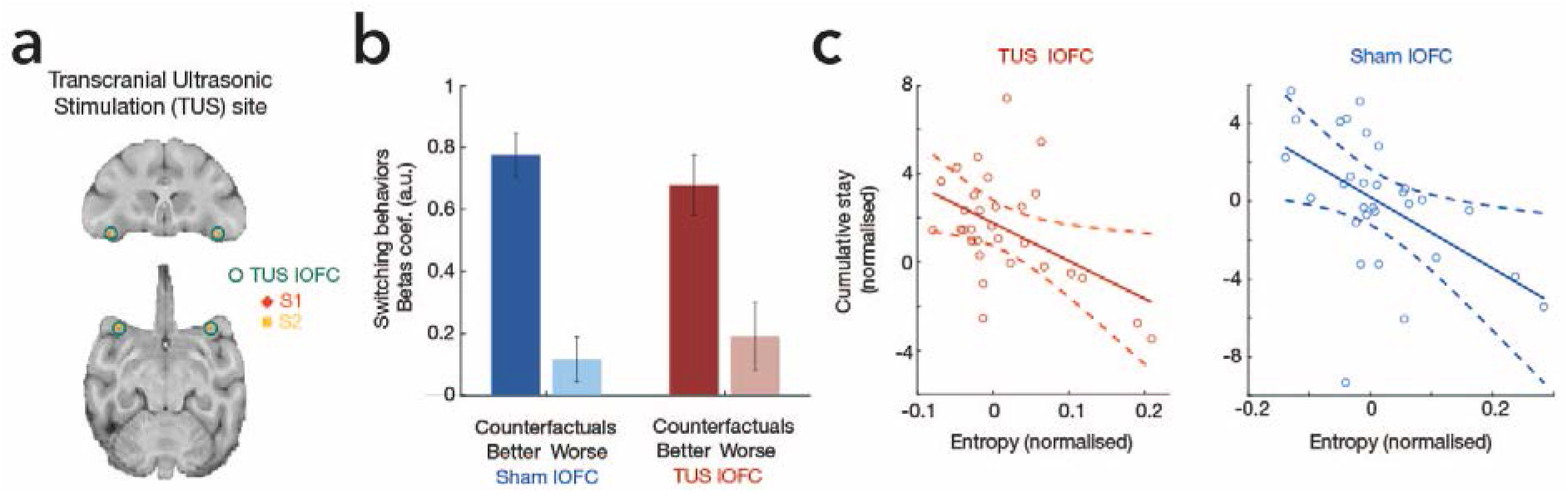
Transcranial Ultrasonic Stimulation of lateral orbitofrontal cortex (lOFC) did *not* impair translation of counterfactual choice values into actual behavioral change. **a)** Neurostimulation site where TUS was applied for each animal (S1 and S2). The TUS transducer was set at a resonance frequency of 250 kHz and concentrated ultrasound in cigar-shaped focal spot in lOFC. **(b)** The significant difference between the influence of the better and worse counterfactual option value on future switching behavior (in blue, as per fig.2f and 5c) was unaltered after lOFC TUS (in red). **(c)** Entropy is strongly and negatively predictive of change in exploratory behavior (indexed by cumulative number of “stay” choices: choices of the same option on one trial after another) in the control condition (blue) and this remains the case after lOFC TUS (red).

**Supplementary Figure 5:**
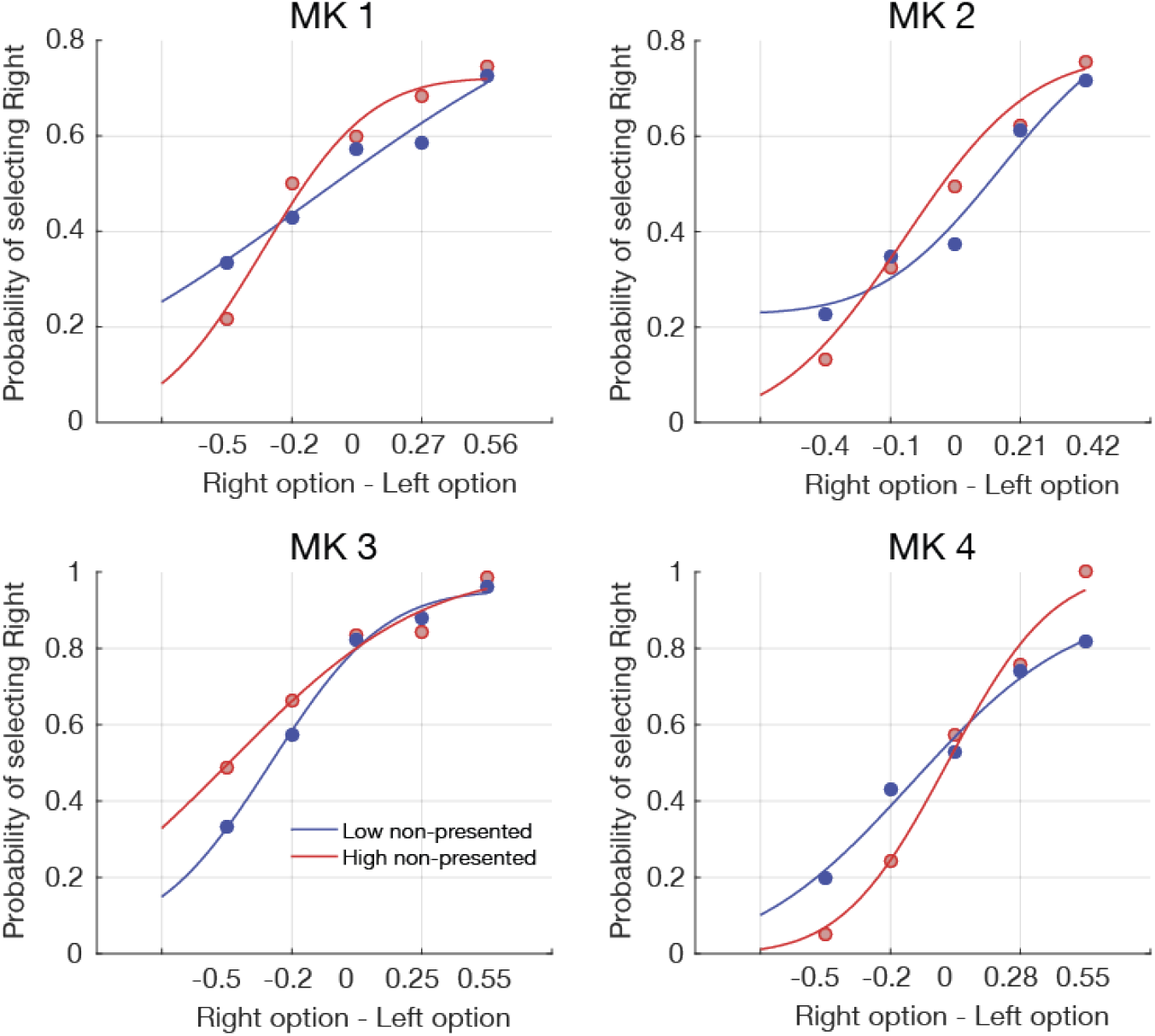
The unavailable option value affects the efficiency of choice behavior. **(a)** Average choice behavior when choosing between the Left and Right options plotted as a function of the value of the unavailable option (low value: blue; high value: red) value for each animal separately (data are averaged across sessions). Curves plot logistic functions fit to the choice data.

## Methods

### Subjects

Four male rhesus monkeys (Macaca mulatta) were involved in the experiment. They weighed 10.4–11.9 kg and were 7 years of age. They were group housed and kept on a 12 hr light dark cycle, with access to water 12–16 hr on testing days and with free water access on non-testing days. All procedures were conducted under licenses from the United Kingdom (UK) Home Office in accordance with the UK The Animals (Scientific Procedures) Act 1986 and with the European Union guidelines (EU Directive 2010/63/EU).

### Behavioral Training

Prior to the data acquisition, all animals were trained to work in an MRI compatible chair in a sphinx position that was placed inside a custom mock scanner simulating the MRI scanning environment. They were trained to use custom-made infra-red touch sensors to respond to abstract symbols presented on a screen and learned the probabilistic nature of the task until reaching a learning criterion. The animals underwent aseptic surgery to implant an MRI compatible head post (Rogue Research, Mtl, CA). After a recovery period of at least 4 weeks, the animals were trained to perform the task inside the actual MRI scanner under head fixation. The imaging data acquisition started once they performed at more than 70% accuracy (choosing the option with the highest expected value) for at least another three consecutive sessions in the scanner.

### Experimental task

Animals had to choose repeatedly between different stimuli that were novel in each testing session (Figure 1a). We used a probabilistic reward-based learning task inspired from tasks originally designed to study reinforcement learning. However, our task consisted of a series of choices, on each trial, between two stimuli drawn out of a larger pool of three and this manipulation alters the nature of reinforcement learning in at least two major ways. First, the subjects have to maintain in memory the value of the option that is not directly available. Second, it creates a horizon of choices that is not deterministic, as the animal cannot predict what option will be presented next. The position of the two available options on the left and right side of the screen were fully randomized. Animals had to choose any symbols by touching one of two infra-red sensors placed in front of their two hands corresponding to the stimuli on the screen. After making their decision, if the correct option was selected, the unselected option disappeared and the chosen option remained on the screen and a juice reward was delivered. If an incorrect choice was made, no juice was delivered. The outcome phase lasted 1.5 seconds. Each reward was composed of two 0.6 ml drops of blackcurrant juice delivered by a spout placed near the animal’s mouth during scanning. Each animal performed six to seven sessions in the MRI scanner. The experiment was controlled by Presentation software (Neurobehavioral Systems Inc., Albany, CA).

### Reinforcement learning algorithms

We used two reinforcement-learning algorithms (Maintain and Decay models) to estimate trial-by-trial expected values associated with each option using animals’ responses ^1^. For both models, if stimulus *A* was selected on trial *i*, its value was updated via a prediction error, *δ*, as follows: *ν_A_*(*i* + 1) = *ν_A_*(*i*) + *α. δ*(*i*) where *a* is the learning rate and the prediction error was given by *δ*(*i*) = *r*(*i*) – *ν_A_*(*i*). The values of the unselected stimulus (e.g. *B*) were not updated. The two models differ in their assumptions of the stimulus that was not shown on that trial (e.g. *C*). In the Maintain model, the values of C were maintained at their current values such that *ν_c_*(*i* + 1) = *ν_c_*(*i*). In the Decay model, the values of C were updated as followed: *ν_c_*(*i* + 1) = *ν_c_*(*i*) + *γ*·(*ν_c_*(1) – *ν_c_*(*i*)). To generate choices for both models, we first used a softmax procedure in which, on every trial, the probability of choosing stimulus *A* was given by: *PA*(*i*) = *σ*(*β*(*ν_A_*(*i*) – *ν_B_*(*i*))) where *σ*(*z*) = 1/(1 + *e^−z^*) is the logistic function, and *β*the degree of stochasticity in making the decision. The model choice probabilities were then fitted against the discrete behavioral choices to estimate the free parameters (*α*, *β*, *γ*).

### Model fitting

To estimate the free parameters (*α, β, γ*), we used separate fitting procedures. The first fitting procedure employed maximum likelihood estimation and a constrained non-linear optimization procedure (as implemented in *fmincon* in MATLAB) separately for each session. The associated likelihood function was given by: 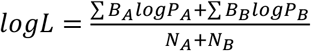 where N_A_ and N_B_ denote, the number of trials in which stimulus A and B were chosen, and B_A_ (*B_b_*) equals 1 if A (*B*) was chosen on that trial, and 0 otherwise. We fitted this function similarly for the other two stimulus combinations (*AC* and *BC*) and found the optimal parameters by minimizing the sum of the three negative log-likelihoods.

The second fitting procedure we employed minimizes overfitting ^2^, and for each session i we found the maximum *a posteriori* estimate of each model’s free parameters: 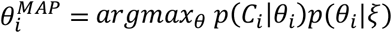 where *p*(*C_i_*\*θ_i_*) is the cross-entropy loss function between observed and predicted choices *C_i_* given the model parameters *θ_i_* and *p*(*θ_i_|ξ*) is the prior distribution on the model parameters *θ_i_* given the population-level hyperparameters ξ. In order to estimate the optimal ξ we implemented an Expectation-Maximization algorithm, which performs k iterations of a two-stage optimization routine until convergence. Particularly, during the Expectation step we optimized the session-wise joint distribution over the data and parameters with respect to the parameters holding the hyperparameters fixed: 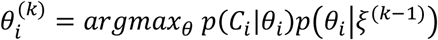 and found the posterior distribution over the parameters using a Laplace approximation: 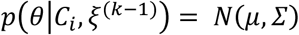. During the Maximization step we revised the population-level hyperparameters ξ by updating the first and second moments of the multivariate normal distribution over the parameters.

### Reinforcement-learning model comparison

To determine the best fitting model we subsequently performed two types of formal Bayesian model comparison amongst the fitted models. The first approach treats each model as a random-effect and is therefore more robust to outliers than fixed-effect approaches ^3,4^. Specifically, we first estimated the session-wise Laplace approximated log evidence for each model. We subsequently computed the model-wise exceedance probability (that is, how confident we are that a model is more likely than any other model tested) using the spm_BMS routine (http://www.fil.ion.ucl.ac.uk/spm/software/). The second approach measures each model’s goodness of fit based on the model’s population-level hyperparameters. We implemented the procedure described by Huys and colleagues ^2^. Here the model log likelihood is obtained by integrating over the model’s parameters. Sampling the model’s parameters from a Gaussian prior density whose mean and variance are set to the population-level hyperparameters allow us to approximate the integral.

### Imaging Data Acquisition

Awake-animals were head-fixed in a sphinx position in an MRI-compatible chair. We collected fMRI using a 3T MRI scanner and a four-channel phased array receive coil in conjunction with a radial transmission coil (Windmiller Kolster Scientific Fresno, CA). FMRI data were acquired using a gradient-echo T2* echo planar imaging (EPI) sequence with 1.5 - 1.5 - 1.5 mm^3^ resolution, repetition time (TR) = 2.28 s, Echo Time (TE) = 30 ms, flip angle = 90, and reference images for artifact corrections were also collected. Proton-density-weighted images using a gradient-refocused echo (GRE) sequence (TR = 10 ms, TE = 2.52 ms, flip angle = 25) were acquired as reference for body motion artifact correction. T1-weighted MP-RAGE images (0.5 - 0.5 - 0.5 mm^3^ resolution, TR = 2,5 ms, TE = 4.01 ms) were acquired in separate anesthetized scanning sessions.

### fMRI data preprocessing

FMRI data were corrected for body motion artefacts by an offline-SENSE reconstruction method ^5^ (Offline_SENSE GUI, Windmiller Kolster Scientific, Fresno, CA). The images were aligned to an EPI reference image slice-by-slice to account for body motion and then aligned to each animal’s structural volume to account for static field distortion ^6^ (Align_EPI GUI and Align_Anatomy GUI, Windmiller Kolster Scientific, Fresno, CA). The aligned data were processed with high-pass temporal filtering (3-dB cutoff of 100s) and Gaussian spatial smoothing (full-width half maximum of 3mm). The data that were already registered to each subject’s structural space were then registered to the CARET macaque F99 template^7^ using affine transformation.

### fMRI data analysis

We employed a univariate approach within the general linear model (GLM) framework to perform whole-brain statistical analyses of functional data as implemented in the FMRIB Software Library ^8^:

*Y = Xβ + ε = β*_1_*X*_1_ + *β*_2_*X*_2_ + … + *β_N_X_N_ + ε* where Y is a T×1 (T time samples) column vector containing the times series data for a given voxel, and X is a T × N (N regressors) design matrix with columns representing each of the psychological regressors convolved with a hemodynamic response function specific for monkey brains ^9,10^. β is a N × 1 column vector of regression coefficients and ε a T × 1 column vector of residual error terms. Using this framework we initially performed a first-level fixed effects analysis to process each individual experimental run which were then combined in a second-level mixed-effects analysis (FLAME 1 + 2) treating session as a random effects (we also had a similar number of sessions across subjects). Time series statistical analysis was carried out using FMRIB’s improved linear model with local autocorrelation correction. Applying this framework, we performed the GLMs highlighted below.

### GLM1 – correct vs. incorrect future selection of the currently unavailable option

Our first fMRI analysis was designed to reveal the brain regions representing the value of the currently unavailable option to guide accurate future decision making. Specifically, locked to the decision time, we included 3 boxcar regressors with a duration of 100 ms that were then convolved with the hemodynamic response function: 1) an unmodulated regressor indexing the occurrence of a decision (Dec; all event amplitudes set to one), 2-3) two parametric regressors whose event amplitudes were modulated by the expected value of the unavailable option for i) future correct selection (unavcorr) and ii) future incorrect selection (unavincorr). Additionally, we included two fully parametric regressors whose event amplitudes were modulated by the expected value of the chosen (Ch) and unchosen (Unch) options that were available on the current trial. Locked to feedback time we included a binary regressor representing positive and negative feedback (+1/-1) and a categorical regressor representing right and left responses (+1/-1), such as:

*Y = β*_1_Dec + *β*_2*unav*_*cor*__ + *β*_3*unav*_*cor*__ + *β*_4_*Ch* + *β*_5_*Unch* + *β*_6_*Fbk* + *β*_7_*Side* + *ε*. Finally, to further reduce variance and noise in the BOLD signal, we add two unconvolved regressors locked at time of feedback and with a duration of a TR (2.28sec) for left and right responses (to capture variance in the BOLD signal caused by any field distortion coincident with responding), six nuisance regressors one for each of the motion parameters (three rotations and three translations), and extra single-trial nuisance covariates for abrupt changes in the BOLD signal.

### GLM2 – Subjective choice comparison (Chosen option value vs. Unchosen option value)

Our second fMRI analysis was designed to reveal the brain regions representing the decision variable guiding choices between the options actually available on the current trial (Chosen option value-Unchosen option value). Locked to decision time, we included 4 boxcar regressors with a duration of 100 ms that were then convolved with the hemodynamic response function: 1) an unmodulated regressor indexing the occurrence of a decision (Dec), 2-4) three fully parametric regressors whose event amplitudes were modulated by the expected value of the chosen option (Ch), unchosen option (Unch) and unavailable option (Unav) and the same covariates of non-interest as described in GLM1:

*Y = β*_1_*Dec + β*_2_*Ch + β*_3_*Unch + β*_4_*Unav + β*_6_*Fbk + β*_7_*Side + ε*. In the third GLM (GLM3: counterfactual model), the Unchosen and Unavailable options were replaced by the Better and the Worse alternatives and in the fourth GLM (GLM4: difficulty model), the Chosen and Unchosen options were replaced by the High Value option and the Low Value option presented.

### Neural model comparison

To assess goodness of fit at the neural level and avoid double dipping in favor of the hypothesis that we wanted to support (GLM3)^11^, we first defined from GLM2, several ROIs within a network including all the brain areas that survived cluster level P < 0.001 (cluster-based correction) for the value comparison (chosen-unchosen) contrast. Within this network, we derived the log-evidence from GLM2, GLM3 and GLM4. Log evidence was then fed into a Bayesian model selection random effects analysis (using the spm_BMS routine), which computed the exceedance probability of each GLM for each ROI. This analysis indicates which GLM best explained the neural data. We report the results for ACC, lPFC, and vmPFC/mOFC.

### ROI analyses

We conducted analyses on ROIs defined as two-voxel radius spherical masks placed over the hippocampus (Right: x = 16.5, y = -7.5, z = -12; left: x = -14, y = -9, z = -12.5 CARET macaque F99 coordinates), ACC (x = 1, y = 20.5, z = 10.5), lPFC (x = 14.5, y = 17.5, z = 9.5), vmPFC/mOFC (x = -5, y = 14, z = 2). We used procedures now standardly employed in most human and animal neuroimaging studies^12–14^ in which the mean and standard error (denoted in all figures by lines and shadings respectively) of all the within-subject b weights were calculated across sessions for plotting the effect size time courses (each animal had a similar number of sessions).

### Leave one out for ROI spatial peak selection

We used a leave-one-out procedure to identify ROI peak voxels for the analyses of main effects for areas identified in all fMRI analyses. For each group level analyses, our procedure involved leaving one session out at a time. From the results of the remaining 24 sessions, we extracted local maxima of the relevant clusters and centered the ROIs for the left out session on the local maxima. We repeated this for all sessions. Therefore, the ROI selection was statistically independent from the data of the session that was subsequently analyzed in the ROI.

### Leave one out on time-series group peak signal

We performed significance testing on time course analyses using a leave-one-out procedure on the group peak signal to avoid potential temporal selection biases. For every session, we calculated the time course of the group mean beta weights of the relevant regressor based on the remaining 24 sessions. We then identified the (positive or negative) group peak of the regressor of interest within the analysis window of 1 to 6 seconds from decision onset. Then, we took the beta weight of the remaining subject at the time of the group peak. We repeated this for all subjects. Therefore, the resulting 25 “peak” beta weights were selected independently from the time course of the subject analyzed. We assessed significance using t-tests on the resulting beta weights.

### Transcranial Focused Ultrasound Stimulation (TUS)

A single element ultrasound transducer (H115-MR, diameter 64 mm, Sonic Concept, Bothell, WA, USA) with a 51.74 mm focal depth was used with a coupling cone filled with degassed water and sealed with a latex membrane (Durex). The ultrasound wave frequency was set to the 250 kHz resonance frequency and 30 ms bursts of ultrasound were generated every 100 ms with a digital function generator (Handyscope HS5, TiePie engineering, Sneek, The Netherlands). Overall the stimulation lasted for 40 s. A 75-Watt amplifier (75A250A, Amplifier Research, Souderton, PA) was used to deliver the required power to the transducer. A TiePie probe connected to an oscilloscope was used to monitor the voltage delivered. The recorded peak-to-peak voltage was constant throughout the stimulation session. Voltage values per session ranged from 128 to 136V and corresponded to a peak negative pressure of 1.152 to 1.292MP respectively measured in water with an in house heterodyne interferometer (see ^15^ for more details about the simulation protocol). Based on a mean 66% transmission through the skull ^16^, the estimated peak negative pressures applied ranged from 0.76 to 0.85 MPa at the target in the brain.

The transducer was positioned with the help of a Brainsight neuronavigating system (Rogue Research, Montreal, CA) so that the focal spot would be centered on the targeted brain region, namely the rACC (F99 coordinates x = 1, y = 20.5, z = 10.5) (identified according to coordinates of the maximum peak used in GLM2). The ultrasound transducer / coupling cone montage was directly positioned to previously shaved skin on which conductive gel (SignaGel Electrode; Parker Laboratories Inc.) had been applied. The coupling cone filled with water and gel was used to ensure ultrasonic coupling between the transducer and the animal’s head.

A sham TUS condition (SHAM) was also implemented as a non-stimulation control. Sham sessions were interleaved with TUS sonication days and completely mirrored a typical stimulation session (setting, stimulation procedure, neuro-navigation, targeting of ACC, transducer preparation and timing of its application to the shaved skin on the head of the animal) except that sonication was not triggered.

### Entropy analyses

For the analyses presented in Fig. 5 (behavioral analysis of TUS data), we used a running window analysis with entropy defined as: 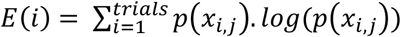, in which x_i,j_ is the probability that a given option j is associated with a positive feedback on trial i. We then used the slope of entropy (difference between the beginning and the end of a window of 20 trials) as a measure of environmental predictability. A positive change in entropy reflects that the environment is less and less predictable and should trigger exploration whereas a negative change in entropy should engage exploitative behavior. As a proxy for exploration/exploitation, we used the cumulative sum of stay behavior, which is simply a vector, keeping track of the number of times a choice has been chosen. Note that a consecutive stay for an option A that has been chosen on trial t could also include trials for which on the next trial (t+1) A would not be available but chosen on the subsequent trial (t+2).

### vmPFC partial regression analysis

To test the strength of the link between the unavailable option’s impact on the current decision and its neural impact in vmPFC/mOFC, we computed the accuracy residuals (Y*, from regressing accuracy against the values of the two available options omitting the unavailable one) and the unavailable residuals (X* from regressing the unavailable option value against the values of the two observable options) and then regressed Y* against X* ^17^ for each session separately (see average effect on Fig.6c).

## Supplementary methods

### Functional connectivity analyses

Psycho-physiological interaction (PPI) analyses were performed in two stages. We first performed a whole brain PPI analysis. Using the ROI procedure described above, we extracted time-series data from individual clusters in the right hippocampus that served as a seed region (that is the physiological regressor: PHY). This analysis was primarily designed to investigate the potential interaction of the hippocampus with decision-related regions involved in selecting the option that had previously been unavailable. As such, the increase in correlation between the hippocampus and potential decision making regions should be specific for the task in which this coupling is relevant; that is, it should be greater during a future trial in which the previously unavailable option is now correctly selected. Therefore our psychological (PSY) task regressor was constructed such that correct selection trials were weighted +1 and incorrect selection trials were weighted −1. The PPI analysis thus included the following regressors during the decision phase of the trial in which the previously unobserved option was now selected: 1) an unmodulated regressor indexing the occurrence of a decision (Dec), 2) the PHY regressor, 3) the PSY regressor and 4) the interaction regressor (PHY × PSY). The rest of the design was identical to GLM 1/2/3.

In a second stage and in order to further confirm that the PPI result actually reflects a negative correlation between hippocampal activity and the vmPFC/mOFC during the correct selection of a previously unavailable option ^18^, we extracted the BOLD time course of the vmPFC/mOFC according to the same procedure described in the ROI section and computed the PPI regressor by taking the vmPFC/mOFC time course and the PSY contrast described above. Note that we did not perform any further statistical analysis on this PPI regression as the significance was already assessed in the whole brain analysis ^11^.

